# resPAINT: Accelerating volumetric super-resolution localisation microscopy by active control of probe emission

**DOI:** 10.1101/2022.04.14.488333

**Authors:** Edward W. Sanders, Alexander R. Carr, Ezra Bruggeman, Markus Koerbel, Sarah I. Benaissa, Robert F. Donat, Ana Mafalda Santos, James McColl, Kevin O’Holleran, David Klenerman, Simon J. Davis, Steven F. Lee, Aleks Ponjavic

**Affiliations:** Department of Chemistry, University of Cambridge, Cambridge, CB2 1EW, UK; Radcliffe Department of Medicine, John Radcliffe Hospital, University of Oxford, OX3 9DS Oxford, UK; Cambridge Advanced Imaging Centre, University of Cambridge, CB2 3DY Cambridge, UK; School of Physics and Astronomy, University of Leeds, Woodhouse Lane, Leeds LS2 9JT, UK; School of Food Science and Nutrition, University of Leeds, Woodhouse Lane, Leeds LS2 9JT, UK

**Keywords:** PAINT, Super-resolution imaging, Volumetric imaging

## Abstract

Points for accumulation in nanoscale topography (PAINT) allows the acquisition of practically unlimited measurements in localisation microscopy. However, PAINT is inherently limited by unwanted background fluorescence at high probe concentrations, especially in large depth-of-field volumetric imaging techniques. Here we present reservoir-PAINT (resPAINT), in which we combine PAINT with active control of probe photophysics. In resPAINT, a ‘reservoir’ of non-fluorescent activatable probes accumulate on the target, which makes it possible to drastically improve the localisation rate (by up to 50-fold) compared to conventional PAINT, without any compromise in contrast. By combining resPAINT with large depth-of-field microscopy, we demonstrate volumetric super-resolution imaging of entire cell surfaces. We then generalise the approach by implementing multiple switching strategies, including photoactivation and spontaneous blinking. We also implement alternative volumetric imaging modalities including the double-helix pointspread function, the tetrapod point-spread function and singlemolecule light field microscopy. Finally, we show that resPAINT can be used with a Fab to image membrane proteins, effectively extending the operating regime of conventional PAINT to encompass a larger range of biological interactions.

## Introduction

Single-molecule localisation microscopy (SMLM) enables routine imaging of biological structures down to a spatial resolution of tens of nanometres.^1^ Fundamentally, biology occurs in three dimensions and therefore there has been an increasing focus on methods that can interrogate biological phenomena over increasingly larger volumes. This has motivated the development of large depth-of-field (DOF) single-molecule imaging techniques, such as the doublehelix point-spread function (DHPSF),^2,3^ the tetrapod pointspread function (PSF)^4^ and single-molecule light field microscopy (SMLFM).^5^ In SMLM, it is becoming necessary to collect substantial numbers of localisations (>100k μm^-3^) to accurately represent increasingly large structures. Obtaining appropriate (*i*.*e*. Nyquist) sampling is difficult because 3D SMLM is limited by a combination of speed of acquisition^6^ (*i*.*e*. the localisation rate), high background in large DOF imaging and photobleaching.^7^ The development of techniques capable of high localisation densities with high contrast and elevated localisation rates would therefore enhance volumetric imaging. This would offer valuable insight into numerous biological questions, including the spatial distribution of biomolecules (*e*.*g*. T-cell activation^8^ and chromatin organisation^9^) as well as the study of protein-protein interactions.^10,11^ To achieve this, it is necessary to design labelling approaches that: 1) minimise fluorescence background and 2) accommodate high emitter densities.

Photoactivation localisation microscopy (PALM^12,13^, Fig. 1a) and direct stochastic optical reconstruction microscopy (dSTORM^14–16^, Fig. 1a) have been used extensively to image biological structures in cells, but these techniques suffer from irreversible photobleaching, which reduces the localisation rate over time. This is particularly problematic for volumetric imaging, where the axial dimension greatly increases the number of localisations required for sufficient sampling.^6^ Probes can be refreshed, but this requires complex microfluidic approaches.^17^ PAINT (Fig. 1a) circumvents photobleaching by the intermittent binding of fluorescent probes to targets from solution. Once probes stochastically bind, they can be localised and subsequently vacate the site or photobleach^18^, thus enabling probe replacement and practically unlimited acquisition times. There are generally two strategies employed: firstly, in conventional PAINT, a fluorescent binder intermittently attaches to the target of interest (*e*.*g*. antibody fragments (Fabs), peptides, antigens, small molecules), but these are often limited by the binding kinetics.^19–23^ A second approach, DNA-PAINT, involves attaching a DNA docking strand to the target, typically *via* immunolabeling or a HaloTag-linker, which facilitates PAINT imaging using transiently binding complementary single-stranded DNA probes in solution.^24,25^

**Fig. 1.**
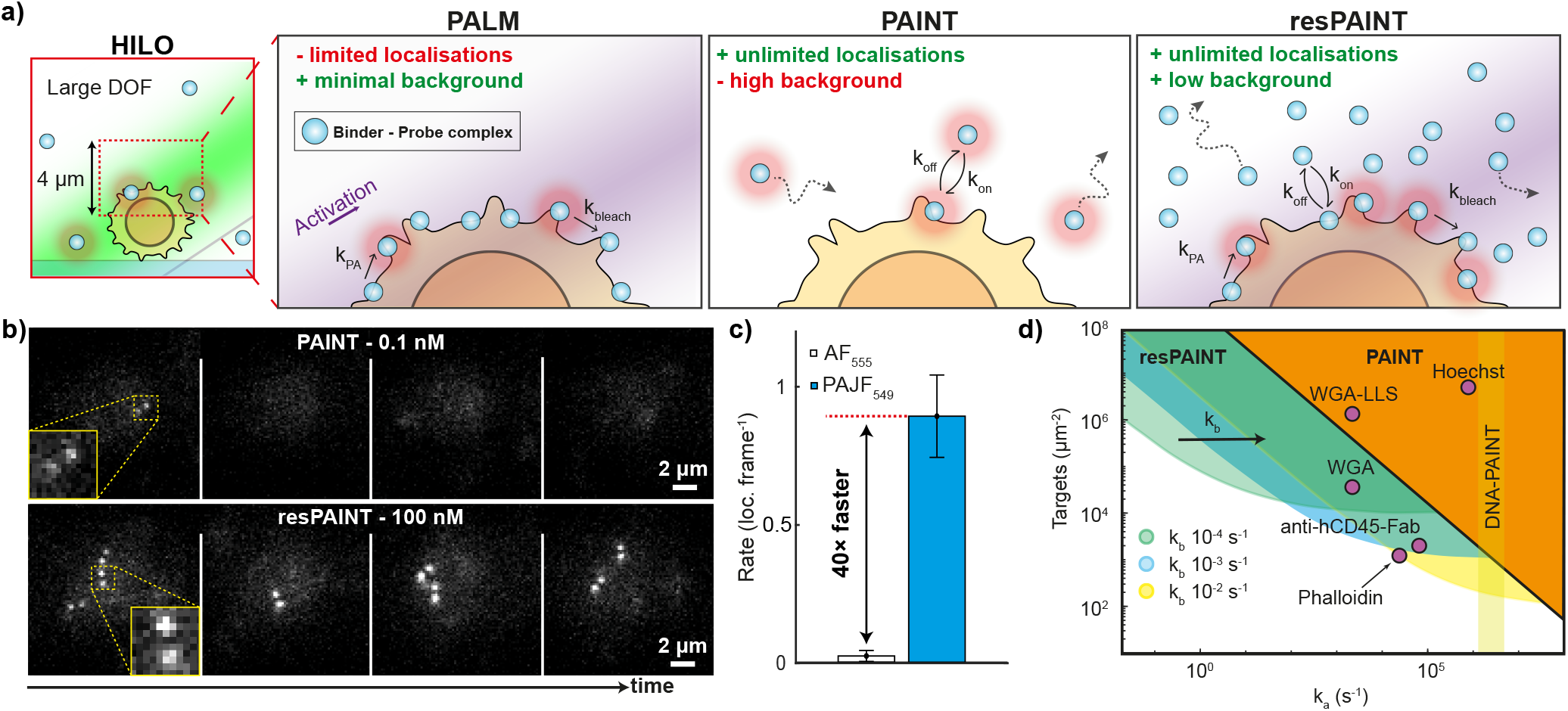
resPAINT combines active control of probe emission with PAINT to greatly enhance localisation rates. **a) HILO:** 3D SMLM is often performed using highly inclined and laminated optical sheet (HILO) excitation as this allows some optical sectioning. When combined with PSF engineering, as in DHPSF, this enables imaging of a large DOF of up to 4 μm, which is useful for imaging complex topography such as the cell surface. **PALM:** PALM uses photoactivation of stably bound probes for SMLM, which achieves high contrast but finite localisation numbers. **PAINT:** In PAINT, probes transiently bind to targets, achieving virtually infinite localisations with the drawback of increased background from unbound probes in solution. **resPAINT:** With resPAINT, we combine active control of probe emission with PAINT to achieve both practically unlimited localisations and high contrast. With most probes being in the non-fluorescent state, much higher probe concentrations can be used to effectively concentrate probes on targets that can be activated to improve localisation rates, without increasing the background. **b)** Representative SMLM time-series taken on the apical surface of fixed Jurkat T cells using conventional PAINT with WGA-AF555 at 0.1 nM and resPAINT with WGA-PAJF549 at 100 nM, demonstrating how the localisation rate is improved for similar backgrounds. **c)** Quantification of (b) showing 40-fold improvement in localisation rate. *n* = 5 cells for each condition. Error bars indicate s.d. **d)** Operational regimes of PAINT and resPAINT with DHPSF imaging for a variety of targets (see main text and Supplementary Note 1 for details).

A major advantage of DNA-PAINT is the ability to tune binding kinetics for optimal SMLM. Despite this, large DOF 3D imaging still presents a significant challenge due to increased background from excess fluorescent probes in solution. This can be prevented by lowering the probe concentration; however, this typically results in prohibitively low localisation rates. Instead, methods have been developed where the probe ‘lights up’ or is concentrated on target *versus* in solution by way of Förster resonance energy transfer (FRET),^26^ fluorogenic probes,^27^ repeat PAINT,^28^ quenching,^29^ photoactivation^30^ and many others. While these strategies greatly improve volumetric imaging with DNA PAINT, they retain inherent drawbacks including: unspecific binding^31^, photocleavage^32^, the necessity of immunolabeling and an inability to observe direct binding events such as peptide/receptor interactions. Application of similar lightup strategies to Fab or small molecule binders would greatly increase the range of biological interactions that can be observed using PAINT. A notable example (where DNA-PAINT cannot currently be used) is actin-staining using the smallmolecule binder phalloidin, which often relies on refreshment of probes from the imaging volume, even in a dSTORM mode.^33^

In PAINT, the background scales linearly with the concentration of the probe *[A]*, while the localisation rate, *dB*_*reservoir*_*/dt*,^34^ is effectively controlled by the concentration of probe, *[A]*, the association rate of the binder, *k*_*a*_, as well as the number of bound sites, *B*_*bound*_, and the total number of the targets, *B*_*max*_ (see Supplementary Note 1 for definitions and discussion of our modelling). Since the association rate is generally fixed for a given binder, the localisation rate, which scales as *k*_*a*_*[A]*, cannot be increased without increasing *[A]*. This inevitably leads to unacceptable levels of background fluorescence (from diffusing probes), as the background also scales linearly with *[A]*. If the target density is high (*i*.*e. B*_*max*_ is large), as for lipid membranes, glycocalyx and nuclear staining, then proteins and small molecule binders may achieve suitable localisation rates.^6,18,20,35–37^ However, in cases of relative molecular sparsity (*i*.*e. B*_*max*_ is small), as for membrane proteins and receptors, the localisation rate is often unsuitable. It is this physical limitation in the binder kinetics that dictates the localisation rate, and ultimately the speed of PAINT.

To solve this issue, we introduce reservoir-PAINT (resPAINT, Fig. 1a), which addresses these issues by using active control of probe emission. By ensuring that most of the probes remain in a non-fluorescent state, the probe concentration can be increased by orders of magnitude without introducing additional unwanted background signal (Supplementary Fig. 1). This results in artificial concentration of the probe on target to create a ‘reservoir’ of bound and non-fluorescent probes that can be activated, and importantly replenished. Note that the accumulation and replenishment of this reservoir depends on the kinetics of a given binder. Therefore, the association rate, *k*_*a*_, and the dissociation rate, *k*_*b*_, dictates the types of biological interactions that can be studied with the technique. Interestingly, there is some suggestion that resPAINT may have been utilised in several previous studies, without formalising or fully realising the concept, where it is typically referred to as no-wash labelling.^33,38–41^ We demonstrate that resPAINT can improve the localisation rate, and therefore the speed of acquisition, for a given binder up to 50-fold. We implement resPAINT using the DHPSF^2,3^ a common volumetric imaging technique. This makes it suitable for capturing large variations in structure such as the complex morphology of the cell surface, which we illustrate by imaging the entire membrane of a T cell. Furthermore, we show that resPAINT is a generalisable principle that works across a variety of probe control mechanisms (photoactivation or pH-tuned spontaneously blinking probes), as well as large DOF volumetric imaging modalities (DHPSF, tetrapod PSF^4^ and SMLFM^5^). Finally, we use the greatly improved contrast to achieve 3D-PAINT imaging of a T-cell membrane protein, CD45, using an antibody fragment, (Fab). The ability to enhance the localisation rate without increasing background greatly extends the application range of PAINT, improving accessibility to volumetric imaging of biological structures and enables super-resolution imaging of a greater variety of biologically relevant ligand-receptor interactions.

## Results

### resPAINT enhances the localisation rate of protein binders

To demonstrate the concept of resPAINT, we first evaluated a standard PAINT probe-binder complex without active control of emission. We used the probe Alexa Fluor 555 (AF_555_) covalently linked to the binder wheat germ agglutinin (WGA), a commonly used cell-membrane stain that binds to the large number of N-glycosyl sites on the cell surface.^6^ First we imaged the apical surface of fixed Jurkat T cells using the DHPSF (Fig. 1b 30 ms exposure, ∼10 kW cm^-2^ power density). We systematically varied the amount of binder to determine the maximum concentration (0.1 nM) that maintained an acceptable background (<16 photons/pixel, see Supplementary Note 1). Under these conditions, the localisation rate was prohibitively small (0.02 loc. frame^-1^, Fig. 1c) and would require ∼50 hours to acquire a dataset with 100,000 localisations to approach Nyquist sampling (50 nm resolution, ∼1000 loc. μm^-2^ for a 2D membrane, imaged in 3D).

Next, we investigated the performance of a photoactivatable (PA) Janelia Fluor probe (PAJF_549_)^42^ attached to WGA for resPAINT. Successful implementation of this concept requires tuning both binder concentration and the photoswitching kinetics, such that a ‘reservoir’ of photoactivable probes can be established on the membrane, *i*.*e*. the binder is artificially concentrated at the target site. Conceptually, if the binder concentration or the photoactivation rate is relatively low, the localisation rate would be impractical for super-resolution imaging. Conversely, if the concentration or photoactivation rate is excessive, the large fluorescence background would decrease the signal to noise ratio, which would deteriorate both the localisation rate and the localisation precision. We imaged fixed Jurkat T cells stained with WGA-PAJF_549_ under identical conditions to that of WGA-AF_555_ but now with photoactivation (power density of 2 W cm^-2^) and a higher probe concentration of 100 nM (1000-fold larger, Fig. 1b). We observed a 40-fold improvement in localisation rate (0.85 loc. frame^-1^) for comparable background levels compared to WGA-AF_555_ (Fig. 1c Supplementary Fig. 1, Supplementary Movie 1). This demonstrates how replacement of a conventional PAINT probe with a photoactivatable probe greatly improves the localisation rate, given suitable binding kinetics. The precision achieved is appropriate for 3D SMLM (18 nm lateral, 38 nm axial, Supplementary Fig. 2a). The constant localisation rate over time (Supplementary Fig. 3-4) confirmed that the probe-binder complex was undergoing PAINT. The large improvement in localisation rate can be rationalised by considering a 1000-fold increase in the binder concentration results in the effective concentration of the probe-binder complex being high, while the observable fraction of the probe remains low (Supplementary Fig. 1).

To explore the experimental regimes of resPAINT, we modelled the photophysical and binding kinetics of activatable probe-binder complexes binding to static targets on a cell (see Supplementary Note 1 for details). Since the localisation rate in PAINT is dependent both on the kinetics of the binder as well as the number of targets, we evaluated typical ranges of target densities and association/dissociation rates to estimate the operating regime of PAINT and resPAINT using the DHPSF (Fig. 1d Supplementary Note 1). The working range was defined as achieving a localisation rate of >1 s^-1^μm^-2^ and a background signal of <0.6 probes μm^-2^. We found that resPAINT greatly extends the range of practical association rates and target densities to a regime where it now becomes possible to use low affinity ligands to observe a protein (or other molecules of interest) using PAINT (Fig. 1d). The suitable range of *k*_*b*_ spanned approximately 10^−2^-10^−4^ s^-1^, covering a variety of biologically relevant interactions.^43–45^ These data show that at lower target densities, it is necessary to have a faster dissociation rate, although it should be noted that *k*_*b*_ can be tuned for a given binder.^46^ A notable example that now becomes accessible with resPAINT, as shown in Fig. 1d is membrane-protein imaging using a Fab.

While we have focused on DHPSF imaging here, resPAINT could be applied to any SMLM modality, such as total internal reflectance fluorescence (TIRF) or confocal-PAINT microscopy.^47^ As shown by our conventional PAINT example, WGA is typically not a suitable binder when using DHPSF, but WGA has been shown to work with PAINT previously (WGA-LLS in Fig. 1d).^6^ However, this required the use of a complex experimental setup, including cyclic imaging and lattice light-sheet microscopy (LLS), to achieve the required background reduction through optical sectioning. In contrast, the enhanced localisation rate afforded by resPAINT allowed us to perform WGA imaging with DHPSF using a comparatively simple setup. Importantly, given suitable binding kinetics, any PAINT application using standard fluorophores could be improved by changing to an activatable probe as in resPAINT.

### Whole-cell volumetric super-resolution imaging using resPAINT

A motivation for developing resPAINT was to enable facile whole-cell volumetric super-resolution imaging of the cell surface. To achieve this, it was necessary to optimise the conditions of our WGA-PAJF_549_ binder-probe complex. The conditions shown in Fig.1b-c (100 nM and ∼2 W cm^-2^ 405 nm power density) were determined by exploring a wide range of concentrations and photoactivation rates to find optimal imaging conditions for PAJF_549_ (Fig. 2a-b, Supporting Movie 2). It should be noted that these conditions will vary depending on the choice of fluorophore and imaging modality. The optimised concentration and activation power densities are specific to WGA, PAJF_549_ and DHPSF imaging. To apply resPAINT with PAJF_549_ to a different system, the conditions determined here should provide a practical starting point for protocol optimisation, provided that the dissociation rate, *k*_*b*_, allows the build-up of a reservoir of bound non-fluorescent probes.

**Fig. 2.**
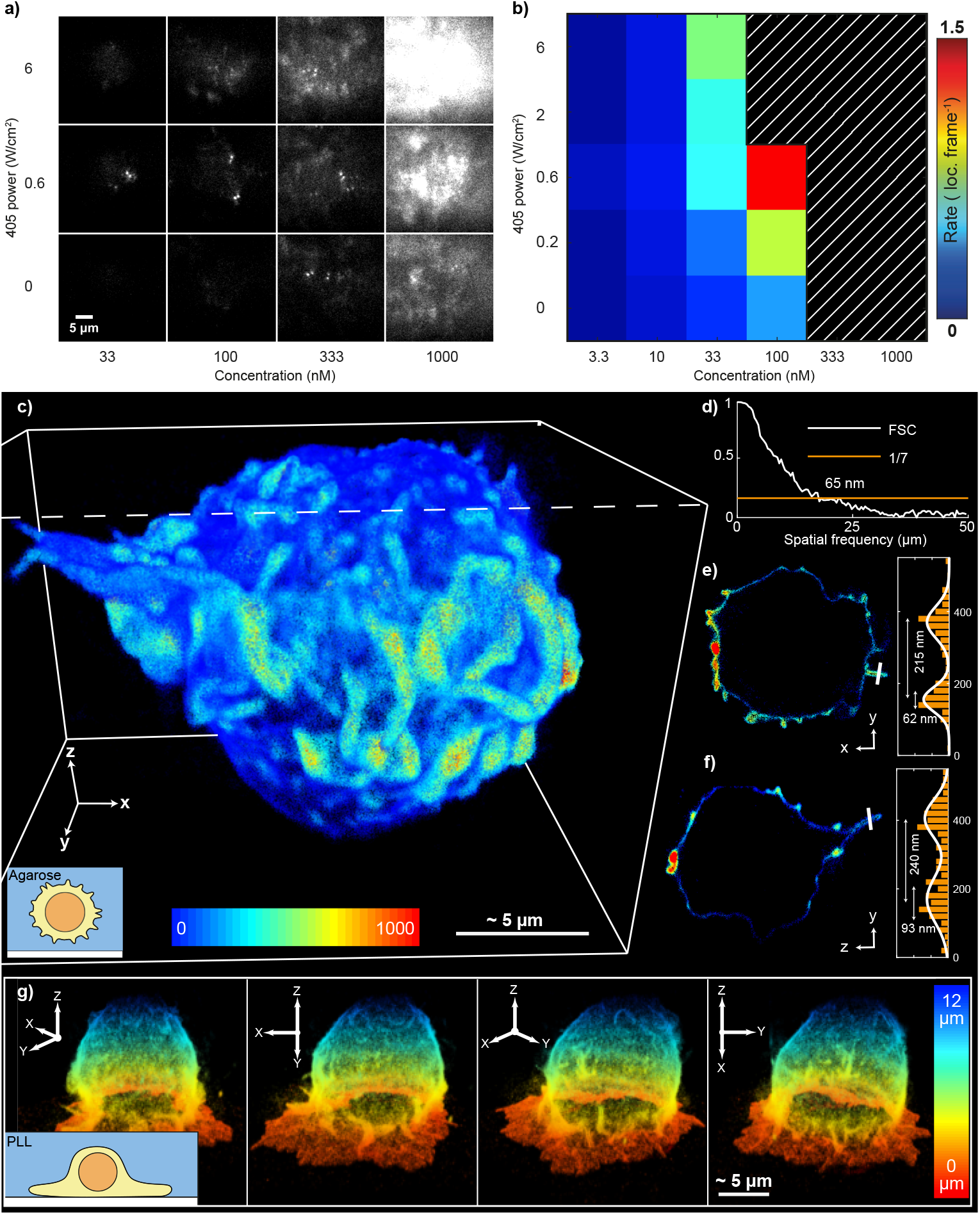
Optimisation of resPAINT to enable whole-cell 3D superresolution imaging of the cell membrane. **a)** Representative resPAINT imaging of the apical surface of fixed Jurkat T cells, using a variety of photoactivation powers and probe concentrations. **b)** Quantification of the localisation rate in (a), highlighting the identified optimal conditions of 100 nM and 0.6 W cm^-2^ activation power. *n* = 5 cells for each condition. The hatched black/white region indicates the area where the background exceeded the threshold of 200 photons. **c)** 3D super-resolution image of the entire membrane of a Jurkat T cell acquired using resPAINT with PAJF_549_ and DHPSF. The image comprises 1,400,000 localisations, collected over 200,000 frames per z-slice at 30 ms exposure time, and was stitched together using four 4 μm z-slices taken in 3.5 μm steps. The cell is coloured by the number of localisations within a local 200 nm radius, to highlight topographical features. The inset depicts a cartoon of how the T cell was suspended in an agarose gel to avoid surface interactions. d) Fourier shell correlation (FSC)^48^ was used to estimate the isotropic resolution in (c), which was found to be 65 nm (1/7 cutoff).e) A y-z slice of the image (c) and corresponding line plot (highlighted in white) through the middle of a microvilli with measured width 240 nm. F) As (e) for a y-x slice, with measured microvilli width of 215 nm. g) As in (c), coloured by height, showing 4 rotated views of a Jurkat T cell that has interacted with a PLL-coated coverslip for 10 minutes prior to fixation. Inset shows a cartoon of a skirt formed due to surface interaction that can be observed in the super-resolution image. A line profile applied to the skirt determines the skirt thickness to be 218 nm.

We have previously used DHPSF to image protein distributions in Jurkat T cells using PA fusion proteins^49^ and dSTORM.^8^ However, the achievable localisation density was always limited by the number of targets on the cell as well as by photobleaching. We applied our optimised WGAPAJF_549_ imaging to whole-cell 3D super-resolution imaging to achieve high localisation density without compromising contrast.

Jurkat T cells were fixed in suspension and then dispersed into an agarose hydrogel matrix containing fiducial markers. Cells were imaged over four axial optical slices (4 μm DOF taken in 3.5 μm steps) to cover the entire cell volume, where adjacent planes shared fiducial markers to enable drift correction and localisation alignment between planes. 800,000 total frames were recorded with a frame rate of 30 ms to acquire 1,400,000 localisations that were used to construct a whole-cell 3D super-resolution image of the T-cell surface (Fig. 2c Supporting Movie 3). We characterised the resolution of the image using Fourier shell correlation (FSC^50^) as 65 nm (Fig. 2d), which compared well to LLS-PAINT^6^ imaging of WGA that demonstrated an estimated FSC resolution of 110 nm. We also assessed the ability of membrane resPAINT to resolve complex morphological features. Line profiles were applied through ‘finger-like’ structures to assess their observable resolution-limited FWHM and membrane thickness (Fig. 2e Fig. 2f), which were found to be 200-250 nm and 60-95 nm respectively in agreement with expectations.^51^

To demonstrate the ability of this approach to capture large-scale deformations in membrane surface topography at high resolution, we also imaged a Jurkat T cell that had been allowed to interact with a poly-L-lysine (PLL)-coated coverslip for ten minutes after which it was fixed (Fig. 2g Supporting Movie 4). This electrostatic coating has been found to induce activation of T cells and associated large-scale morphology changes of the membrane.^52,53^ Here, resPAINT was able to capture the formation of a large ‘skirt-like’ structure (thickness of 218 nm FWHM based on line profile, Supplementary Fig. 5) at the glass-cell interface, demonstrating the dramatic effect of PLL on T cell-surface interactions. In both whole-cell imaging experiments, we observed a constant localisation rate over extended timeframes (∼7 hours) demonstrating that resPAINT is resistant to out-of-focus photobleaching (Supplementary Fig. 4a).

### resPAINT is compatible with multiple activation modes

Next, to demonstrate that the principle of resPAINT can be applied to different classes of probes, and is not limited to photoactivation only, we also studied spontaneously blinking probes (Fig. 3a).^54–56^ This was a good alternative fluorescence switching strategy because: 1) these probes do not require the use of potentially cytotoxic 405 nm laser; and 2) their off-switching rate can be more finely controlled. We implemented spontaneously blinking resPAINT with the probe HMSiR^55^ that undergoes fluorescence intermittency via an intramolecular spirocyclisation reaction (Fig. 3a). The position of the cyclisation equilibrium, *K*_*cyc*_ *= k*_*open*_*/k*_*close*_, is affected by pH, which facilitates control of the ring opening, *k*_*open*_, and ring closing, *k*_*close*_, reactions. We therefore hypothesised that pH could be used to optimise conditions for resPAINT, by shifting the equilibrium, *K*_*cyc*_.^55^ Control of the blinking off rate would also afford greater compatibility across a range of excitation powers and exposure times, compared to the photoactivation implementation. Given a slow dissociation rate, the duty cycle of the probe is now dominated by spontaneous ring closing rather than photobleaching rate.

**Fig. 3.**
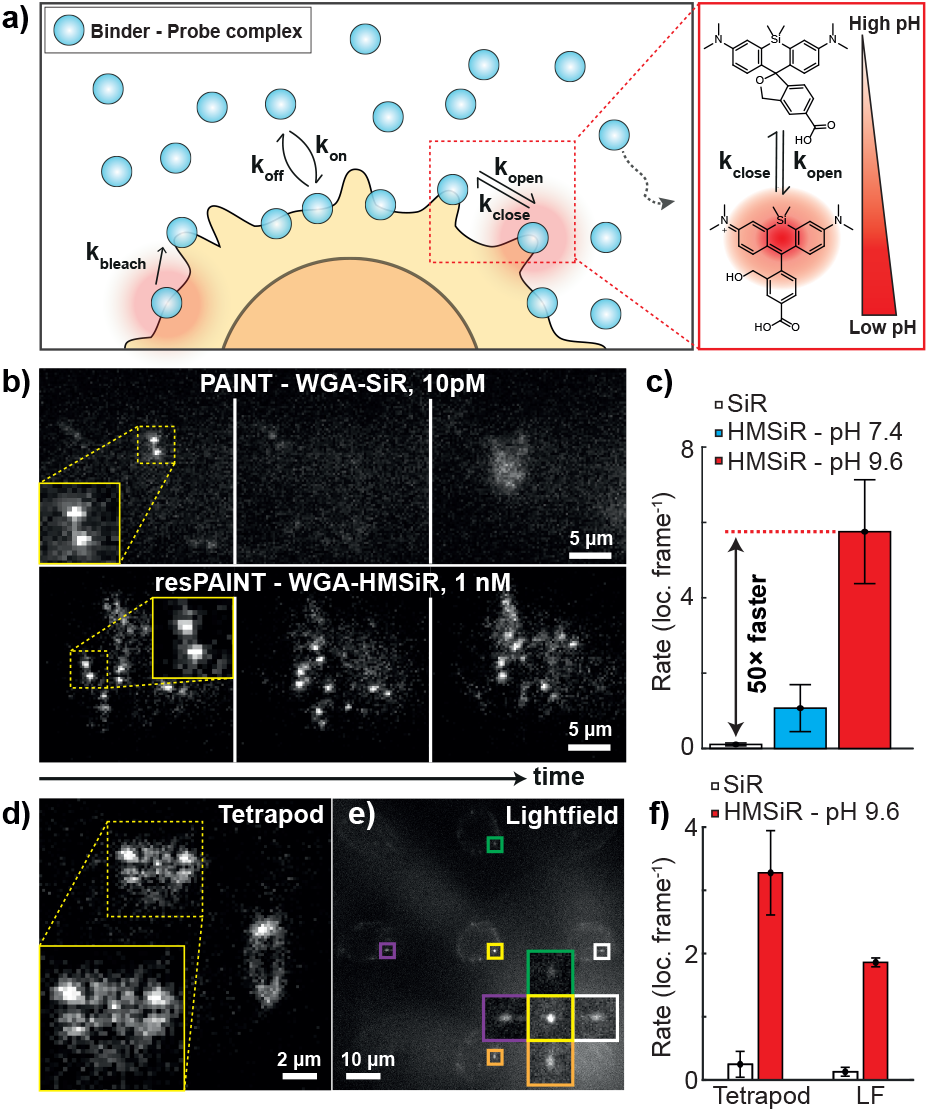
resPAINT using a spontaneously blinking probe. **a)** Left: Cartoon demonstrating the concept of resPAINT using a spontaneously blinking probe. The alternative method of control is achieved via manipulation of the cyclisation equilibrium, K_cyc_ = k_open_/k_close_. Right: A schematic showing the tuning of K_cyc_, visualising the pH dependency of K_cyc_, where at high pH most of the probe is inactive. **b)** Representative SMLM time-series taken on the apical surface of fixed Jurkat T cells using conventional PAINT with WGA-SiR at 10 pM and resPAINT with WGA-HMSiR at 1 nM and pH 9.6, demonstrating how the localisation rate is improved for similar backgrounds. The display contrast was adjusted individually for each condition to aid interpretation. **c)** Quantification of localisation rate under background-matched conditions, demonstrating a 50-fold improvement in localisation rate at pH 9.6. **d-e)** Representative resPAINT images taken on the apical surface of Jurkat T cells using the tetrapod PSF (d) and single-molecule light-field microscopy (e). In (e), the inset shows the same molecule viewed from 5 different angles. **f)** Quantification of (d-e) showing the improvement in localisation rate afforded by resPAINT. *n* = 5 cells for each condition. Error bars indicate s.d.

We demonstrated spontaneously blinking resPAINT using WGA on fixed Jurkat T cells, but now without photoactivation. We first compared the performance of WGA-HMSiR to that of a conventional PAINT probe, WGA-SiR, and observed a modest improvement (10-fold) in localisation rate for a similar background in PBS (pH 7.4) buffer. We found that this pH was suboptimal, owing to an inefficient *k*_*close*_ (effective off time of 245 ms^55^). This timescale was unsuitable for fast imaging (20-100 Hz), while the relatively large on-off ratio limited the effective accumulation of a probe reservoir on the cell. When we increased the pH using a sodium carbonate buffer with pH 9.6 (HMSiR effective off-time of 17 ms^55^), we observed a dramatic improvement in localisation rate (1 nM WGA-HMSiR compared to 10 pM WGA-SiR in PBS) for similar backgrounds (Fig. 3b Supplementary Movie 5). Under these conditions WGA-HMSiR achieved a localisation rate of 5.8 loc. frame^-1^ compared to 0.1 loc. frame^-1^ for WGA-SiR, corresponding to a 50-fold improvement (Fig. 3c). This localisation rate resulted in some overlapping fluorophores and we therefore determined the optimal concentration for WGA DHPSF imaging to be 330 pM. This resulted in a localisation rate of 1.77 loc. frame^-1^ that was stable over one hour (Supplementary Fig. 4b, Supplementary Movie 6), achieving localisation precisions of 22 nm laterally and 50 nm axially (Supplementary Fig. 2b). Increasing the pH further would limit the number of photons collected as the blinking duration would now be shorter than the exposure time (Supplementary Movie 7).

Next, we investigated the performance of resPAINT with alternative extended DOF techniques. We imaged fixed Jurkat T cells using the tetrapod PSF (10 μm DOF,^57^ Fig. 3d) and the recently developed SMLFM (5 μm DOF,^5^ Fig. 3e). Under conditions similar to the DHPSF, we observed improvements in the localisation rate of 13-fold for the tetrapod PSF (Supplementary Movie 8) and 14-fold for SMLFM (Supplementary Movie 9), which was lower than for the DHPSF due to overlapping PSFs (Fig. 3f). Appropriate labelling densities for both tetrapod and SMLFM (Fig. 3d-e) are given in the supplementary information (Supplementary Movies 10 and 11).

### resPAINT imaging using a Fab

Having optimised and applied resPAINT to cell surface imaging, we then evaluated the technique in a more challenging scenario, *i*.*e*., using a Fab to image a membrane protein using PAINT. Conventionally, imaging of low-density targets has necessitated the use of DNA-PAINT with antibodies (Fig. 1d) or chemical alteration of Fab off-rates to enable efficient probe refreshment.^46^ DNA-PAINT typically requires separate imaging and docking strands to form a PAINT pair, which precludes imaging of direct binder-target interactions. Conversely, resPAINT can observe the binder-probe complex and protein target interaction directly in an experimentally straightforward manner (Fig. 4a). We labelled an anti-hCD45Fab (hereafter referred to as ‘Fab’) with either HMSiR or SiR. We then investigated the binding of Fab to protein tyrosine phosphatase CD45 in human Jurkat T cells due to the pivotal role it plays in T-cell activation.^58^

**Fig. 4.**
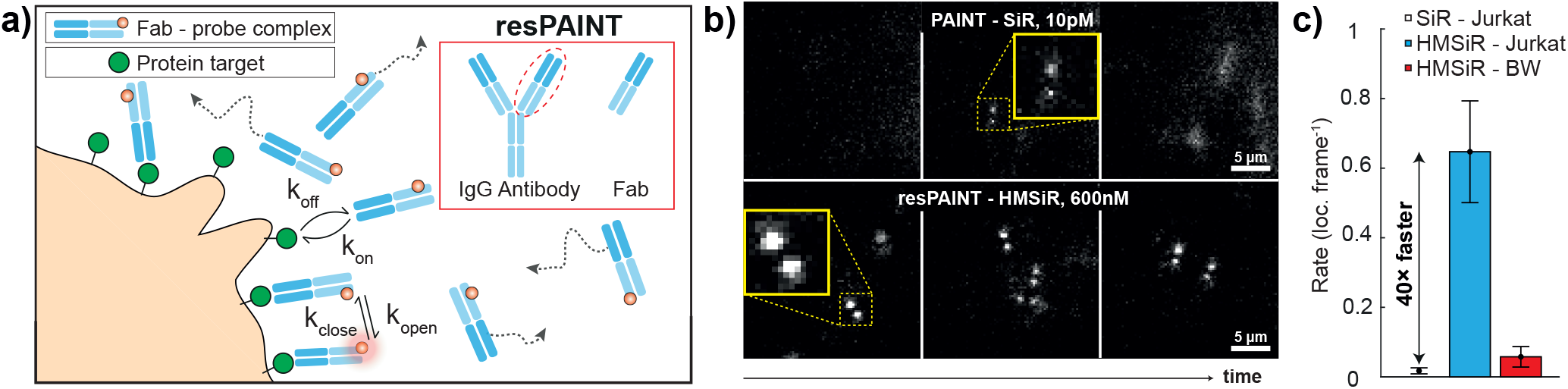
resPAINT with a Fab against CD45. **a)** Cartoon highlighting a resPAINT experiment where a protein is directly imaged using a Fab. Inset, a schematic of a full-antibody and a Fab, where cleavage of the Fab region from the Fc region creates a fragment of the antibody with suitable kinetics for resPAINT. **b)** Representative SMLM time-series taken on the apical surface of fixed Jurkat T cells using conventional PAINT with Fab-SiR at 10 pM and resPAINT with Fab-HMSiR at 600 nM (pH 9.6), demonstrating how the localisation rate is improved for similar backgrounds. **c)** Quantification of the localisation rate as a function for PAINT, resPAINT and a mouse cell control to which the anti-human Fab-HMSiR does not bind mouse-CD45. *n* = 5 cells for each condition. Error bars indicate s.d.

We have previously imaged CD45 on T cells in 3D using dSTORM and the DHPSF.^8^ Therefore, we compared PAINT imaging using 10 pM Fab-SiR with resPAINT using 600 nM Fab-HMSiR (Fig. 4b), which exhibited comparable background levels (Supplementary Movie 12). We first measured the *k*_*b*_ of the Fab bound to fixed Jurkat T cells, which we determined as 1.31×10^−3^ s^-1^ at pH 7.4 and 1.65×10^−3^ s^-1^ at pH 9.6 (Supplementary Fig. 6). These values agree with surface plasmon resonance measurements (1.59×10^−3^ s^-1^, Supplementary Fig. 7) and lie within the previously defined operational regime (Fig. 1d). As in membrane imaging, resPAINT improved the localisation rate 40-fold (0.61 loc. frame^-1^, Fig. 4c) with localisation precision of 22 nm laterally and 56 nm axially (Supplementary Fig. 2c). To confirm the specificity of the Fab binding to human CD45, we used a murine CD45 control cell line, to which the Fab lacks cross-reactivity, and observed a minimal number of localisation events (0.05 loc. frame^-1^, <9 % unspecific binding, Supplementary Movie 12). These results demonstrate that resPAINT increases the accessible range of binder kinetics beyond conventional PAINT.

## Discussion

resPAINT is a method for dramatically improving localisation rates in PAINT without comprising contrast. This is particularly useful for large DOF volumetric imaging, as the technique facilitates acquisition of high localisation densities to achieve Nyquist sampling. The enhancements observed in this work should apply universally, provided the following conditions are met: 1) The target density ranges between 10^3^- 10^8^ μm^-2^ to support the concentrations required for reservoir accumulation and 2) the binder has a *k*_*b*_ where the binding duration permits reservoir build up without inhibiting probe refreshment by saturation of targets. PAINT is also limited by these factors, but resPAINT extends the range of viable binders and target densities. This makes it possible to image targets with relatively low abundance and also alleviates the necessity for high association rates, *k*_*a*_, such that commonly used binders including Fabs, Hoechst and phalloidin now become more accessible for PAINT (Fig. 1d). This allows the use of highly specific antibody-antigen interactions with low affinities in a resPAINT imaging mode, which is typically selected against in the functional characterisation step in traditional monoclonal antibody production.^59^

resPAINT limitations include the requirement to tune the localisation rate via two independent control mechanisms (concentration and switching) which imposes technical complexity. While some aspects of optimisation would be specific to the probe and switching mechanism being used, we demonstrate that activation rates can be controlled via laser power (PAJF_549_) or pH (HMSiR). Importantly, the optimisations performed for these probes would apply to any system in which they are used. Therefore, resPAINT can be applied immediately in other PAINT systems as well as with other probe-target complexes (*e*.*g*. photoactivatable,^42,60^ spontaneously blinking,^54–56^ or fluorescent proteins^61^).

Aspects of the resPAINT principle have been partially explored in previous studies. The combination of photoactivatable probes and collisional flux has been shown to improve super-resolution imaging in materials science with interface PAINT (iPAINT).^62^ However, this study had no bioimaging application, nor did it provide any kinetic framework. Within bioimaging, there is evidence for using switching and collisional flux, although these studies also lack a formal mechanistic description, or indeed may have applied the concept unknowingly. These implementations have typically been referred to as no-wash labelling protocols.^33,38–41^ We provide the first detailed description of the kinetic requirements of resPAINT and explore the space over which the technique is useful for bioimaging. We generalise this concept by using a selection of probes (photoactivation and spontaneously blinking), various imaging modalities (DHPSF, tetrapod PSF, SMLFM) and apply the technique to multiple systems (whole-cell, membrane topography and membrane proteins). The flexibility and extended operational regime of resPAINT demonstrated here makes the technique applicable to numerous biological applications.

The most closely related set of techniques would be the suite of DNA-PAINT tools, where the rapid and tunable binding kinetics of DNA strands make DNA-PAINT highly ubiquitous within SMLM, due to the high localisation precision and compatibility with low target densities. Recent modifications to DNA-PAINT enhance contrast and improve localisation rates in a similar fashion to resPAINT by adopting various ‘light-up’ strategies.^24,26,28–30^ Furthermore, the use of left-handed DNA has improved specificity in DNA containing samples.^31^ When compared directly, resPAINT offers some advantages over DNA-PAINT in that: 1) it is fully compatible with DNA containing samples; 2) conjugation with DNA can be experimentally complex and 3) DNA-PAINT can suffer from binding-site depletion, although appropriate buffers can somewhat mitigate this.^32^ Indeed, a potential application of resPAINT would be imaging DNA in whole cell nuclei with Hoechst. This was recently achieved in 3D using Hoechst-JF_646_ in a stimulated emission depletion (STED) PAINT mode.^63^ Hoechst-HMSiR has previously been used in 2D super-resolution imaging,^38^ although the authors argued that they were not operating in PAINT mode. We suggest that they may have inadvertently been using the resPAINT principle based on established Hoechst binding kinetics, the high concentration employed and no-wash labelling. Exchanging JF_646_ with HMSiR and altering pH, or using a photoactivatable Hoechsts,^64^ may yield further improvements.

We show use of a Fab to localise membrane proteins. The use of Fabs with PAINT (Fab-PAINT) has been achieved by using specialised buffers to tune the binding kinetics but requires TIRF sectioning.^54^ These techniques are not mutually exclusive and a combination of Fab-PAINT with resPAINT may be used to achieve even greater improvements in contrast to enable compatibility with HILO illumination and may not require addition of thiocyanate to the imaging buffer.

## Conclusions

We have demonstrated how resPAINT achieves an up to 50-fold improvement in contrast or localisation rate by concentrating probes on target. This in turn extends the operational regime of conventional PAINT and facilitates volumetric 3D super-resolution imaging using large DOF techniques. By simply switching to a probe with active control, it becomes possible to improve existing implementations of PAINT, as long as there is an excess of targets that benefit from an increase in the effective concentration. We hope that resPAINT will simplify and enable future volumetric SMLM applications in previously inaccessible areas that could include: actin PAINT imaging using phalloidin,^33^ intracellular LIVE-PAINT with fluorescent proteins,^22,23^ pPAINT with signalling proteins^10^ and IRIS with peptide fragments.^35^

## ACKNOWLEDGEMENTS

S.F.L. and A.P. conceptualised, supervised and administered the project. E.S., A.R.C. and A.P. developed the labelling methodology, conducted experiments and analysed DHPSF data. E.B. assisted in collection and analysis of tetrapod PSF data. K.O. assisted in the collection of light field data, and S.I.B. processed the data. M.K. assisted with collection of preliminary Fab imaging experiments, and collection of Fab off rate data on cells, while R.F.D. performed surface plasmon resonance experiments. A.M.S., J.M., D.K. and S.J.D provided strategic input on experimental design and provided cell samples. A.M.S and S.J.D. provided anti-hCD45-Fab proteins. E.S., S.F.L. and A.P. wrote the manuscript. All authors reviewed and edited the manuscript.

## AUTHOR CONTRIBUTIONS

The authors would like to thank Dr. C. Leterrier as well as Prof W.E. Moerner and his lab for useful discussions. This work was supported by programme grant funding from the EPSRC (EP/R513180/1, to E.S.), Wellcome trust (206291/Z/17/Z) and programme grant funding from the Royal Society (EP/M003663/1 to A.C). The work was also funded by a Royal Society University Research Fellowship (UF120277, to S.F.L., RGF\EA\181021 For E.B), and by a University of Leeds University Academic Fellowship, as well as an AMS Springboard Award (SBF006\1138) awarded to AP.

## Methods

### Cell culture

Jurkat T cells (ATCC TIB-152) were grown in RPMI (Sigma Aldrich, Madison, WI) while mouse thymoma BW5147 cells were cultured in Joklik-modified Minimum Essential Medium (JMEM (Sigma Aldrich). Both culture media were supplemented with 10% fetal calf serum (FCS) (PAA), 10 mM HEPES (Sigma Aldrich), 1 mM sodium pyruvate (Sigma Aldrich), 2 mM L-glutamine and antibiotics [50 units penicillin, 50 μg streptomycin and 100 μg neomycin per mL] (Sigma Aldrich)). Cells were maintained at 37 °C and 5 % CO_2_ during culturing and, typically, kept at a density between 5-9 × 10^5^ cells mL^-1^.

### Protein Labelling

The corresponding protein WGA (L9640, Sigma-Aldrich, UK) or αCD45 Fab (prepared as in previous work)^8^ was added in a 1:10 molar ratio to the desired dye - PAJF_549_ (Tocris Bioscience, UK) or HM-SiR (Sarafluor-650B, Kishida Chemical Company, Japan) - in 0.02 μm filtered (6808-2002, Cytiva, MA) phosphate buffered saline (PBS, 10010-023, Gibco, MA). 1 M sodium bicarbonate (Sigma-Aldrich) in ultrapure water was added to achieve a 0.1 M concentration in the reaction volume and the reaction was left in the dark for 1.5 hours at room temperature. The protein-dye conjugate was purified by 3 rounds of size exclusion chromatography (Bio-Spin 6 column, BioRad, CA) and then aliquoted in 5 μL portions and stored at -80C until required.

### Cell preparation

∼10^6^ of cells were centrifuged (600g, 2 minutes) and the supernatant was removed before washing once with filtered PBS. The cells were fixed in 0.8 % paraformaldehyde (28906, Thermo Scientific, MA) and 0.2 % glutaraldehyde (G5882, Sigma-Aldrich) for 15 minutes at room temperature. The cells were then washed three times in filtered PBS and then resuspended in ∼100 μL of filtered PBS.

### Cell-coated coverslip preparation for apical surface imaging

Glass slides (24 × 50 mm borosilicate, thickness No. 1, VWR international, PA) were cleaned for 30 minutes with argon plasma (PDC-002, Harrick Plasma, Ithaca, NY) and then coated with poly-L-lysine (PLL, 150-300 kDa; P4832; Sigma-Aldrich) for 15 minutes. The slides were then washed three times with filtered PBS before ∼ 30 μL of PBS was placed with 5-20 μL of cells in PBS and the cells were allowed to settle on the surface for 45 minutes.

### T-cell coated coverslip preparation for whole cell imaging

T cells were adhered to a coverslip using PLL as before. Fiducial markers were prepared as follows. A 100 μL solution containing 50 μm agarose beads (20349, ThermoFisher) was incubated with PLL solution in a 1:1 ratio for 10 minutes followed by centrifugation at 1500 × g for 1 minute. The beads were washed three times with filtered PBS and incubated with nitrogen vacancy fluorescent nanodiamonds (798134, Sigma) in a 1:1 ratio for 10 minutes at room temperature. The labelled beads were then washed three times with filtered PBS and resuspended in 100 μL of filtered PBS. Onto a pre-prepared T-cell coated coverslip was added 3 μL of fluorescent nanodiamond-coated 50 μm diameter agarose beads, which were allowed to settle on the surface. The sample was heated to 37°C and 50 μL of 1 % agarose solution in filtered PBS was added and allowed to settle for 10 minutes. The sample was then allowed to cool to room temperature before 50 μL of filtered PBS was added to the set agarose.

### DHPSF Microscopy

The microscope used for Fab and WGA imaging with the DHPSF was as in our previous work,^49^ incorporating a 1.27 NA 60× water immersion objective lens (1.27 na Plan Apo VC 60×, Nikon) and a quad-band dichroic (Di01-R405/488/561/635-25×36, Semrock) with minor alterations to the optics in the emission path. Namely, for SiR and HMSiR dyes a phase mask (PM) optimised to a different wavelength (650 nm, Double-Helix, Boulder, CO) was used and the fluorescence signal isolated by placement of band-pass and long-pass filter (FF02-675/67-25 and BLP01-647R-25, Semrock) placed immediately before the camera. Excitation light on the sample was filtered using a bandpass filter (FF01-640/14-25, Semrock) In the case of the dyes AF_555_ and PAJF_549_ the PM was replaced with a 580 nm optimised PM (DoubleHelix, Boulder, CO). The fluorescence signal was isolated by use of band-pass and long-pass filters FF01-580/14-25 and BLP02-561R-25 (Semrock, Rochester, NY) and collected by an EMCCD (Evolve Delta 512, Photometrics, Tucson, AZ) operating in frame transfer mode. The excitation light was filtered by use of a bandpass filter (LL02-561-25, Semrock). The DHPSF was calibrated by use of Tetraspeck beads (Thermofisher, T7279) for both PMs and filter combinations, where the fluorescent bead slides were prepared on PLL coated coverslips as in previous work.^49^

### resPAINT imaging of apical T-cell surface

The liquid was carefully removed from a T cell coated coverslip and the surface then gently washed with a prediluted solution of probe at the required concentration in either filtered PBS or in the case of HMSiR in filtered pH 9.6 sodium carbonate-bicarbonate buffer and then imaged on the custom-built DHPSF microscope. For experiments that involved HMSiR and SiR, a 20 ms exposure time was used in both the WGA and Fab imaging cases and a continuous 641 nm excitation beam at (∼5 kW cm^-2^) used in a HILO illumination configuration. The photoactivation mode experiments were conducted with 30 ms exposure times while a continuous 561 nm excitation beam (∼10 kW cm^-2^, measured after objective) was used in combination with a continuous 405 nm beam used for activation at a range of power-densities during optimisation experiments (∼ 0-6 W cm^-2^, measured after objective). An image was collected that centred on the apical surface and contained most of the DHPSFs 4 μm depth of field. In order to quantify the resPAINT improvement, the background was matched in conventional PAINT and resPAINT cases by titrating probe into the imaging volume before an average z-project of an area off cell was taken and the counts measured for a small ROI in the centre of the centre of the frame. The localisation rates at similar background were compared. The quoted improvements are indicative of the difference in localisation rates under these matched conditions.

### Whole cell resPAINT imaging

A coverslip prepared for whole cell experiments was imaged using the same excitation and emission path configuration as for the optimisation PAJF_549_ experiments. Continuous 561 nm illumination (∼ 5kW cm^-2^, measured before objective) and 405 nm excitation (∼ 5 W cm^-2^, measured before objective) was incident on the sample. Four ∼4 μm planes were imaged, where each position contained at least one fiducial marker shared with adjacent planes to allow alignment of localisations post-drift correction. 200,000 frames were recorded at 30 ms exposure for each plane and then the focus was shifted in 3.5 μm steps using a piezo motor. An auto-focus script written in Beanshell was used to maintain the axial position of the sample while acquiring in individual planes.

The resolution of resulting images was evaluated using Fourier shell correlation with a custom MATLAB script. The 3D point cloud dataset was randomly split into two equal parts. This was then used to create a 3D image using 10 nm^3^ voxels, where each point contributed to a Gaussian intensity distribution with σ_xy_ = 40 nm and σ_z_ = 60 nm. Finally, an existing script ^48^ for Fourier shell correlation in MATLAB was applied to the two images to determine the resolution at the 1/7 intercept.

### anti-hCD45-Fab off rate imaging and calculation

T cells were prepared and adhered to a coverslip using PLL as for apical surface imaging before incubation with 200 nM of anti-hCD45-Fab-SiR (Gap8.3-Fab-SiR) for 15 minutes. Imaging was performed on a bespoke microscope as in previous work^66^ using a 641 nm excitation laser (Obis, Coherent). The beam was filtered with an appropriate excitation bandpass filter (FF01-640/14-25, Semrock) and circularly polarised using a wavelength specific quarter-wave plate. The beam was then expanded, collimated and aligned for epifluorescence with an air immersion objective (20× Plan Fluor, NA 0.5, air immersion, Nikon Corporation) mounted on an inverted microscope body (Eclipse Ti2, Nikon Corporation). Emitted light was collected by the same objective lens and separated from excitation light by way of a dichroic mirror (Di01-R405/488/561/635, Semrock) and an appropriate emission bandpass filter (FF01-692/40-25, Semrock). The emitted light was then expanded and focused onto an electron-multiplying charge-coupled device (Evolve 512, Photometrics) for imaging, where the pixel size was 535 nm. A stack of single images was taken in 20 s intervals, with an EM gain of 250, where the exposure time was 100 ms and the power density incident on the sample was ∼0.3 Wcm^-2^. The off rate was measured by fitting the decay in fluorescence signal overtime to an exponential function in Fiji.

### anti-hCD45-Fab surface plasmon resonance measurements

Gap8.3-CD45 interactions were analysed on a Biacore 8k instrument (Cytiva Life Sciences) at a flow rate of 10 μl min^-1^. Running buffer was HBS-P. A Protein A Chip (Cytiva Life Sciences) was used to capture Gap8.3 (∼2000RU) onto Flow cell 2 (FC2) at 10 μl min^-1^. Before injection of CD45 the chip surface was conditioned using 3 injections of HBS-P for 60 s each. Serial dilutions of CD45D1-D4 or CD45RABC were injected for 60 s at 30 μl min^-1^ over both FC1 (reference) and FC2 using single cycle kinetics with a final dissociation time of 300 s. A blank run was also performed using PBS in HBS-P to match the serial dilutions of CD45 for blank subtraction and together with FC1 used for double reference subtraction. All measurements were performed at 20 °C. Results were analysed using the Biacore Evaluation Insight Software (Cytiva Life Sciences) using 1:1 kinetic model binding.

### DHPSF fitting

The whole cell dataset was fitted using easyDHPSF^67^ as previously described. ^49^ Briefly, a calibration dataset was acquired by scanning the stage in 40 nm steps. Using the calibration file, camera parameters and manually selected thresholds, easyDHPSF produced a point cloud of localisations. Drift was corrected based on individual fiducial markers present in each plane. The five planes were aligned by identifying overlapping fiducial markers between planes and correcting localisation positions. Repeated localisations were removed via temporal filters where a localisation was removed if within 500 nm and 0.5 s of a previous localisation. For the images presented and analysed in Fig. 2 a density filter with 200 nm radius was used to remove spurious noisy localisations with less than 5 neighbours.

For all other datasets, DHPSF fitting was done using a custom MATLAB script (currently available at https:github.com/TheLaueLab/DHPSFU). Image sequences were first analysed with the GDSC plugin PeakFit,^68^ to extract localisations. These were paired using the DHPSFU MATLAB script, which uses a PSF calibration file to accurately assign x,y,z positions to the point pairs. Repeat localisations within 20 frames and a 200 nm radius were combined into singles using a temporal filter (∼0.5 s depending on exposure time).

### Tetrapod microscopy

A piezoelectric deformable mirror (DMP40-F01, Thorlabs) was used to generate a tetrapod PSF based on a previous implementation.^57^ The deformable mirror was placed in the conjugate back focal plane of the objective (Plan Apo, 60 ×A/1.40 Oil, DIC H, inf/0.17 WD 0.21, Nikon) using a relay of achromatic doublet lenses (AC254-200-A, Thorlabs). The deformable mirror was controlled using the manufacturers software (version 3.2, Deformable Mirror Software Package, Thorlabs). The tetrapod pattern was generated using a 0.25:-0.75 ratio of secondary and primary astigmatism. The microscope setup was based on a Nikon Eclipse Ti2-E. Two 638 nm diode lasers (each 180 mW, 06-MLD 638 nm, Cobolt) were focused to the back focal plane of the objective using a lens (AC254-250-A, Thorlabs) on a linear translation stage to allow HILO illumination. A dichroic (Di01-R405/488/532/635, Semrock), and emission filters (FF02-675/67-25 and BLP01-647R-25, Semrock) were used. The power density at the sample was ∼2.5 kWcm^-2^. A sCMOS camera was used (Prime 95B, Teledyne Photometrics) and controlled with μManager 2.0 gamma.^69^ Data analysis was performed using the ImageJ plugin ZOLA-3D.^57^ The experimental PSF was modelled using 66 Zernike coefficients. Localisations were filtered based on goodness of fit and photon number (>2000 photons).

### Light field Microscopy

A bespoke lightfield microscope was used as in previous work.^5^ Jurkat T cell membranes were imaged using WGA-SiR or HMSiR with continuous excitation at 638 nm (∼1 kW cm^-2^). HILO illumination configuration was used to minimise fluorescence background to image a plane near the apical surface of Jurkat T cells. Quantification of resPAINT improvement was conducted in the same way as for DHPSF images.

### Light field fitting

The microlens array in the SMLFM system encodes the 3D position of the point emitters in the displacement of the focused image from the optical axis of each lenslet. Sub-diffraction localisation of the point emitter images were performed by fitting a 2D gaussian profile using the ThunderSTORM package.^70^ The 3D localisation was estimated using the previously described method.^5^ This uses knowledge of the optical model and the set of 2D localisations to estimate a 3D localisation for each point emitter. The 3D fitting parameters were: Perspective views (3-5), 2D Gaussian fitting widths (0.4 - 1.2), paraxial angle for grouping (0.5), 3D fit threshold (0.5 μm) and intensity threshold of (200 photons).

