## Supplementary Information for "resPAINT: Accelerating volumetric super-resolution localisation microscopy by active control of probe emission"

### Technical Note 1: Photophysical kinetics of resPAINT

Consider a system where

[A] = concentration of binder-probe complex
$B_{max}$ = total number of binding sites for A
$B_{bound}$ = number of bound sites
$B_{reservior}$ = number of bound activatable molecules
$B_{active}$ = number of bound fluorescent molecules
$B_{PB}$ = number of bound photobleached molecules
$k_a$ = association rate of the binder
$k_b$ = dissociation of the binder
$k_d$ = equilibrium constant
$k_s$ = switching rate of the probe
$k_{PB}$ = photobleaching rate of the probe

$$B_{bound} = B_{reservoir} + B_{active} + B_{PB}$$

$$Number\ of\ vacant\ sites = B_{max} - B_{bound}$$

A binder, labeled with a blinking fluorophore, is added to an aqueous solution at a concentration [A]. The intermittent binding to a target on cells is governed by an association rate, $k_a$, and a dissociation rate, $k_b$. The fluorophore is assumed to be in a dark state given a large on-off ratio. Thus, the concentration of target sites bound with dark fluorophores, $B_{reservoir}$, depends on the association kinetics and the total available binding sites, $B_{max}$.[1] The dark fluorophore can switch into a fluorescent state, via photoactivation or spontaneous blinking, with a switching rate, $k_s$. Assume that the proportion of dye switching from dark to photobleached is negligible. Under these conditions, the change in the dark bound fluorophore concentration, $d(B_{reservoir})/dt$, can be described by

$$\frac{d(B_{reservoir})}{dt} = k_a[A](B_{max} - B_{bound}) - k_b B_{reservoir}$$

where $B_{active}$ is the bound fluorescent concentration and $B_{PB}$ is the bound photobleached concentration. The fluorescent binder can dissociate with the dissociation rate of the binder, $k_b$ or it can photobleach with a rate $k_{PB}$. Note that the on-switching rate of bound dark fluorophores, $k_s B_{reservoir}$, is the quantity that is measured in the experiment and corresponds to the localisation rate in SMLM. In spontaneously blinking probes, the switch off rate is determined by $k_{PB}$ and the rate of ring closing ($k_{close}$ in Fig 3a). For simplicity, we assume that the probe does not undergo spirocyclisation multiple times, *i.e.* the photobleaching rate is dominant. Under these conditions, the change over time, $dt$, in the fluorescent binder concentration, $B_{active}$, can be described by

$$\frac{d(B_{active})}{dt} = k_s B_{reservoir} - k_b B_{active} - k_{PB} B_{active}$$

The photobleached binders can dissociate with the dissociation rate of the binder, $k_b$. Under these conditions, the change in photobleached binder concentration, $B_{PB}$, can be described by

$$\frac{d(B_{PB})}{dt} = k_{PB} B_{active} - k_b B_{PB}$$

Consider the case of WGA at a concentration [A] = 330 nM binding to a cell membrane imaged with DHPSF. In this case $k_a$ = 5,300 $M^{-1}s^{-1}$, $k_b$ = 1.2×$10^{-3}$ $s^{-1}$, equilibrium constant, $k_d$ = 230 nM, and $B_{max}$ = 3.8×$10^6$, assuming 100 μ$m^2$ membrane area imaged with DHPSF at lectin density 3.8×$10^4$ μ$m^{-2}$.[2] WGA is labeled with PAJF$_{549}$ with $k_s$ = $10^{-3}$ $s^{-1}$ and $k_{PB}$ = 100 $s^{-1}$, which depend on activation and excitation power densities.

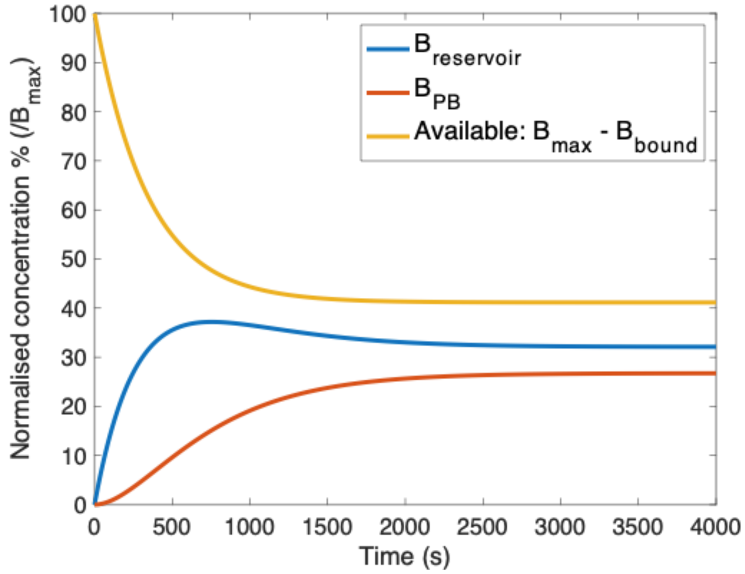


**Figure A1: resPAINT kinetics.** Evolution of bound reservoir, bound photobleached and unbound populations on the cell membrane over time. The reservoir, $B_{reservoir}$, approaches a steady state from which probes can be both replenished and activated for localization imaging.

**Building a reservoir.** Fig. A1 shows how these populations evolve over time. Initially, there is a build-up of bound dark sites that can be activated. As these switch into a fluorescent state and eventually photobleach, there is a build-up of bound photobleached sites. Due to the binding kinetics of WGA, an equilibrium is reached at around 10 minutes, where there is a constant supply of bound dark sites that can be activated for localisation microscopy. As the localisation rate is proportional to $B_{reservoir}$, the maximum is reached in approximately 600 seconds or 10 minutes for WGA-PAJF$_{549}$.

This analysis enables comparison between conventional PAINT and resPAINT for a specific probe. Owing to the fast photobleaching rate, the background is controlled by the activation rate of probes in solution of the observable volume, $V$, in the experiment, given by $k_s[A]V$, while the localisation rate is given by $k_sB_{reservoir}$. In a typical DHPSF imaging volume, $V$, of 10×10×4 μ$m^3$, the background is 80 localisations while the localisation rate is 1,220 loc.$s^{-1}$. PAINT would have a similar background at a concentration of $[A]k_s^{-1}$ = 330 pM, due to fluorescent probes effectively instantaneously diffusing into the excitation volume. At this point the localisation rate in PAINT would be $k_a[A]B_{max}$ = 6.52 loc. $s^{-1}$, or rather resPAINT could reach a theoretical improvement upper limit of up to 188 times faster than PAINT in the case of WGA. Empirically we determine an increase of ~50 fold, this is likely explained by multiple assumptions made in the this analysis (*i.e.* rate constants taken from different cellular systems[2]) and the imperfect nature of the single-molecule experiments (*e.g.* diffusion out of the excitation volume, homogeneity of the excitation volume).

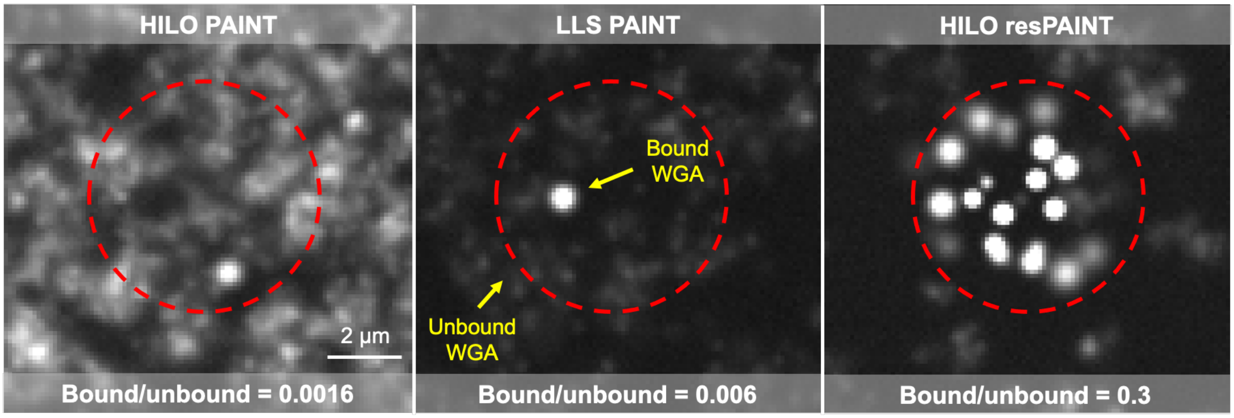


**Figure A2: Simulation of an abundant target (glycocalyx).** Binding and photophysical kinetics for WGA with a conventional PAINT fluorophore (1 nM) and a photoactivatable resPAINT fluorophore (100 nM, $k_s$ = 0.001 $s^{-1}$). The red circle indicates the cell area. The bound/unbound ratio represents the signal/background ratio. LLS improves conventional PAINT 4-fold by reducing out-of-focus excitation, whereas resPAINT provides a 180-fold improvement.

**Simulations.** These findings can be supported by simulating the images that would result from these binding and photophysical kinetics (as described above). We simulated the diffusion ($D$ = 76 $\mu m^2 s^{-1}$) of WGA-probe complexes in solution to compare the performance of PAINT, lattice light-sheet PAINT [3] and resPAINT with similar backgrounds across all three techniques. We evaluate the quality of the images by considering the bound-active/unbound-active molecule ratio, where higher is better.

HILO PAINT with WGA produces poor image quality as there is a large excitation volume, resulting in unwanted fluorescence background caused by emissive diffusing probes. This can be ameliorated somewhat by moving to a confined excitation geometry like LLS. For WGA at 1 nM (Fig. A2, Supplementary Video 13), LLS improves the image 4-fold, to give a suitable background that enables single-molecule imaging at the given localisation rate. When HILO is combined with resPAINT, the excitation volume (and therefore background) is again increased 4-5 times. Despite this, due to building up of the reservoir (through higher concentration 100 nM), resPAINT can still generate substantially, up to 180-fold, better image quality compared to PAINT.

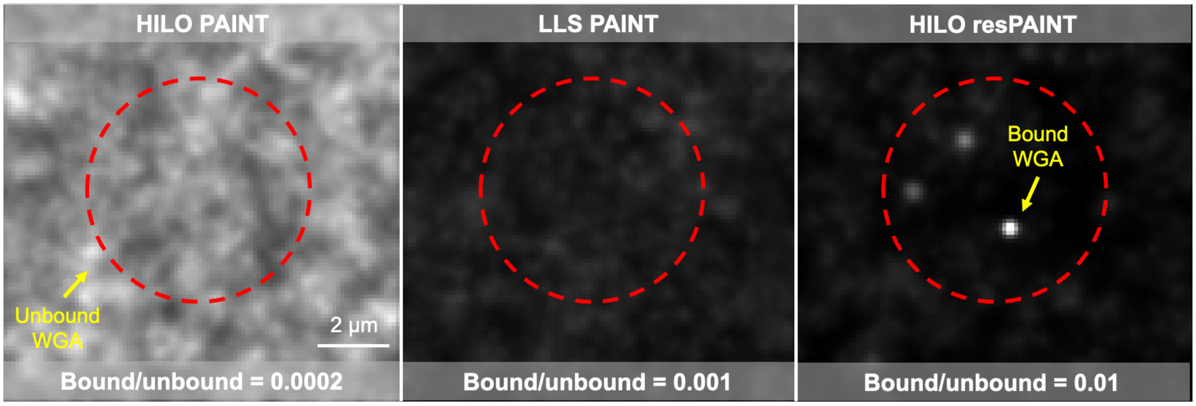


**Figure A3: Simulation of sparse target case** - Binding and photophysical kinetics for a Fab (30,000 targets on cell) with a conventional PAINT fluorophore (10 nM) and a photoactivatable resPAINT fluorophore (1000 nM). resPaint becomes essentially required for imaging under these conditions.

When the target density, $B_{max}$, is reduced as in the case of a protein binder, like a Fab, the binder concentration needs to be greatly increased to provide a suitable localisation rate. In this case resPAINT becomes essential for imaging (Fig. A3, Supplementary Video 14).

**Exploring the accessible regime of resPAINT (Fig. 1d).** By solving the system of linear differential equations, the equilibrium concentration of $B_{reservoir}$ can be determined. This was used to create Fig. 1d. The switching rate, $k_s$, and dissociation rate, $k_b$, were varied over large ranges, for a given association rate, $k_a$, and receptor density, $B_{max}/A$, where $A$ is area. If a condition was found where the localisation rate was larger than 1 $s^{-1}\mu m^{-2}$ and the background was lower than 0.6 molecules $\mu m^{-2}$ (matching LLS conditions[3]) then it was coloured in Fig. 1d for resPAINT and PAINT respectively.

These thresholds were selected as they agreed with our experimental conditions for WGA-PAJF$_{549}$. Therefore, they should be interpreted as a guide for imaging regimes rather than an absolute quantification. However, they should provide a good starting point for identifying whether a certain application would be feasible with resPAINT. Furthermore, the relative scaling of PAINT to resPAINT is linear such that the improvement is independent of the choice of thresholds. Practically this means that for different thresholds, the targets would remain static on Fig. 1d, while the boundaries would move.

### Supplementary Figures

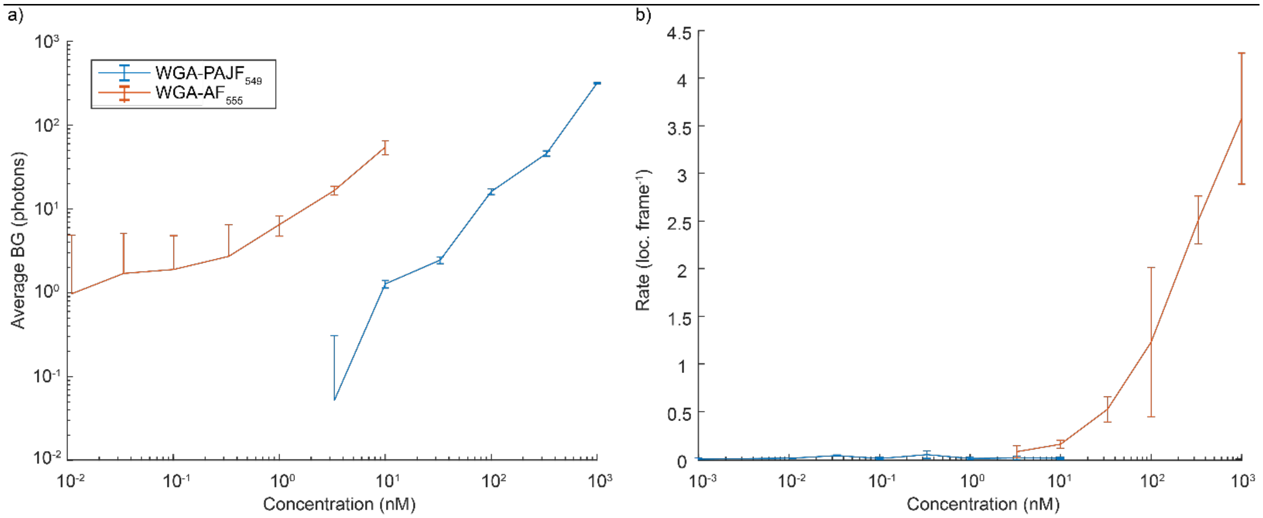


**Supplementary Figure 1: Relating background fluorescence and localisation rate to concentration for both conventional PAINT and resPAINT. a)** Background fluorescence as a function of concentration for both WGA-AF$_{555}$ and WGA-PAJF$_{549}$. In both cases, background increases linearly with concentration. The data show that for comparable levels of background, WGA-PAJF$_{549}$ can support orders of magnitude larger probe concentrations due to the background suppression of resPAINT. *n = 5* cells for each condition. Error bars indicate s.d. **b)** Localisation rate as a function of concentration for WGA-AF$_{555}$ and WGA-PAJF$_{549}$. Owing to the ability to use far higher probe concentrations, and the formation of an inactive probe reservoir on targets, resPAINT can achieve improved localisation rates for comparable levels of background. n = 5 cells for each condition. Error bars indicate 1 standard deviation.

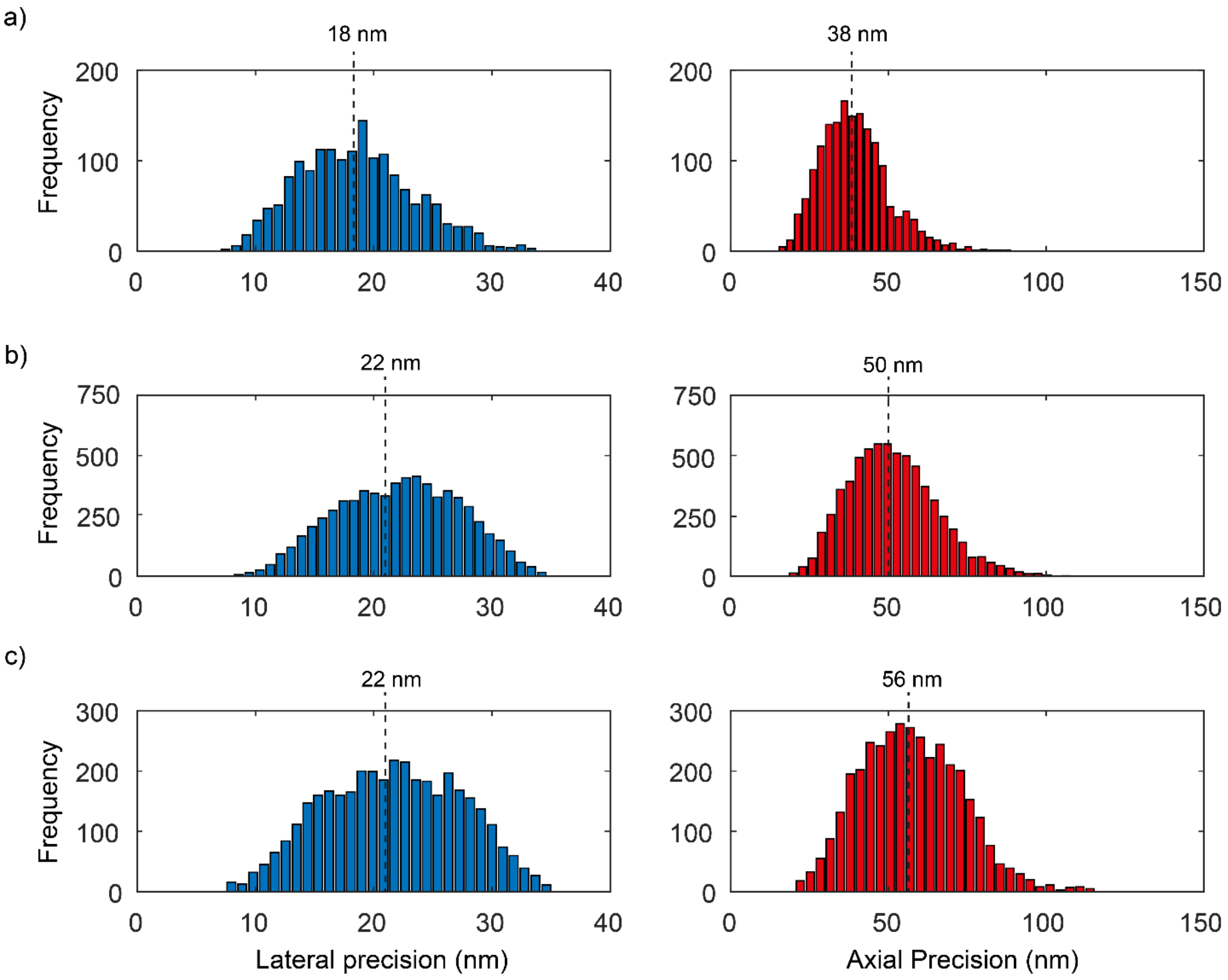


**Supplementary Figure 2: Localisation precision in resPAINT imaging.** Precision histogram for typical resPAINT experiments with **a)** WGA-PAJF$_{549}$ **b)** WGA-HMSiR and **c)** anti-hCD45Fab-HMSiR. We determine median lateral precisions (left) of 18 nm, 22 nm and 22 nm for WGA-PAJF$_{549}$, WGA-HMSiR and anti-hCD45Fab-HMSiR respectively. Median axial precisions (right) were also determined as 38 nm, 50 nm and 56 nm for WGA-PAJF$_{549}$, WGA-HMSiR and anti-hCD45Fab-HMSiR respectively.

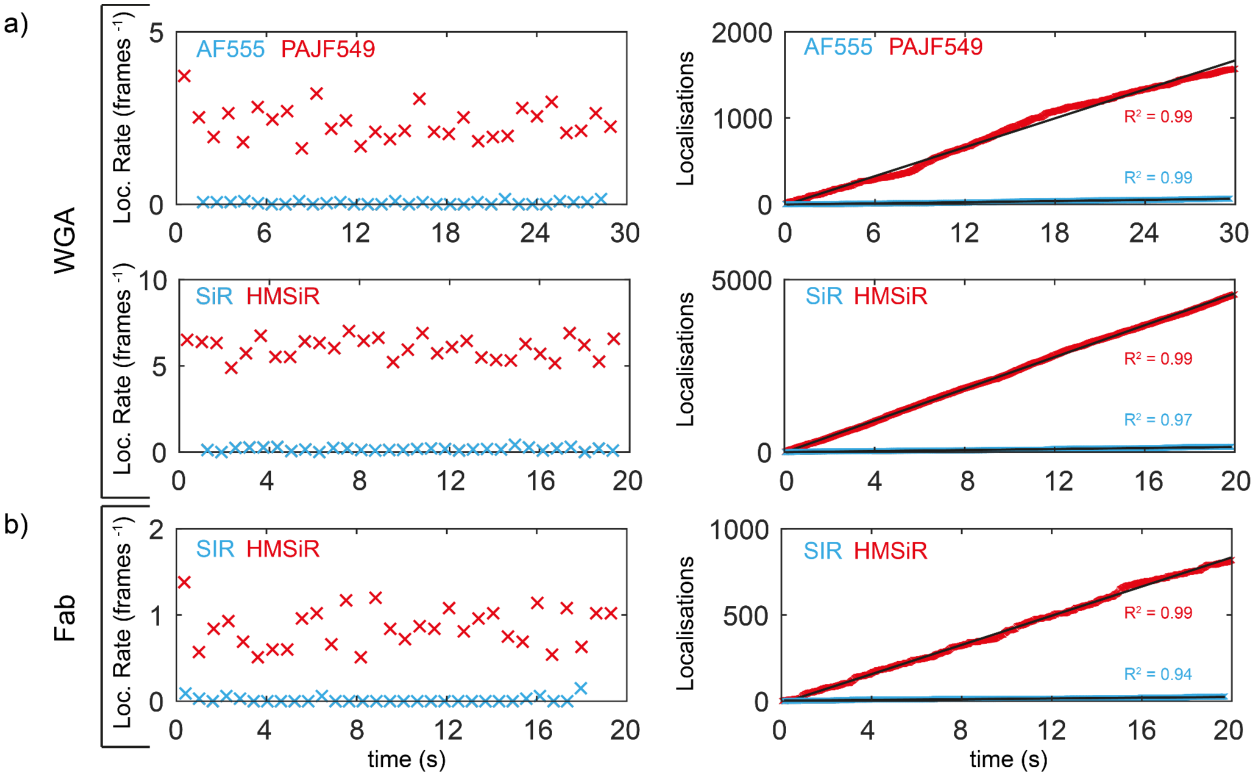


**Supplementary Figure 3**: **resPAINT maintains a stable localisation rate over time, indicative of a PAINT binding mode. a)** Localisation rate and cumulative localisations as a function of time for membrane imaging with WGA **Left:** Histograms of localisation rate as a function of time (bin width is 1 second for PAJF$_{549}$/AF$_{555}$ and 0.6 second for HMSiR/SiR). In both cases, the localisation rate does not change appreciably over time, with resPAINT probes affording a significantly higher localisation rate. **Right:** Cumulative localisations as a function of time, showing a linear response. **b)** As in (a), but for anti-hCD45Fab imaging of CD45 membrane protein and the corresponding conventional PAINT experiment with SiR. A constant localisation rate with time is indicative of PAINT-style imaging.

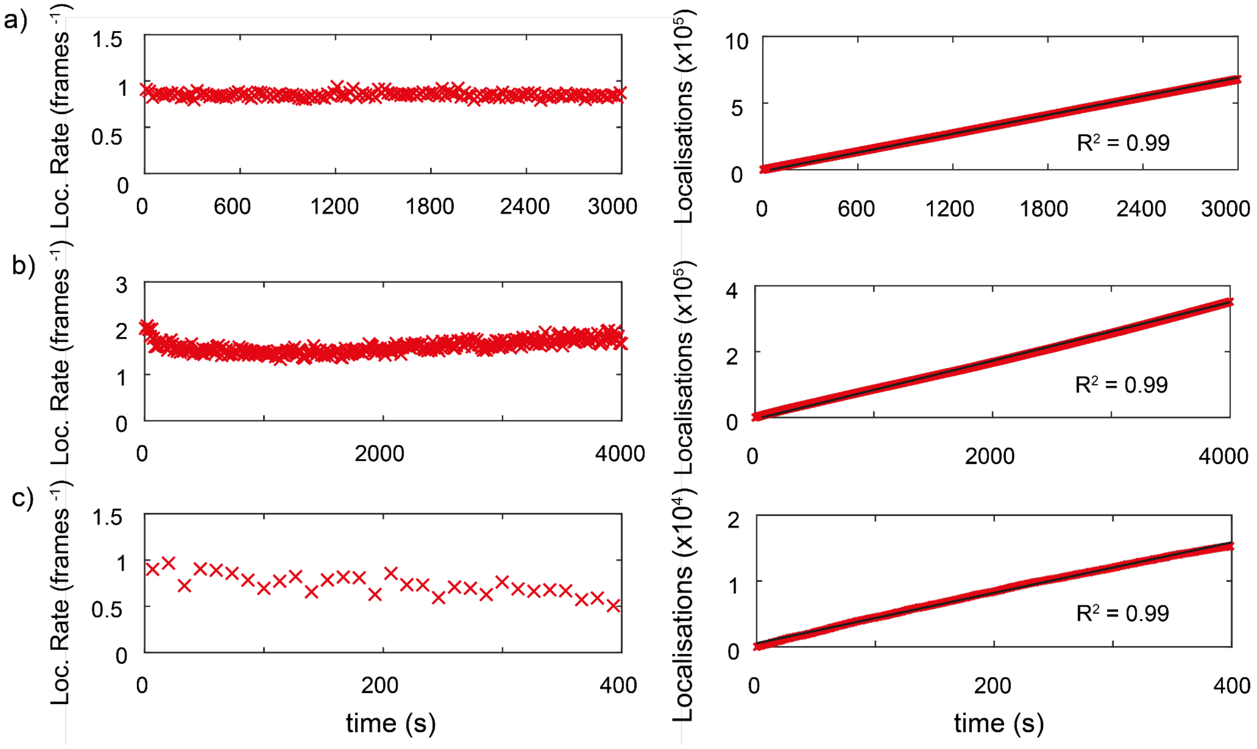


**Supplementary Figure 4: resPAINT maintains a stable localisation rate over long durations indicative of a PAINT binding mode. a)** WGA-PAJF$_{549}$ imaging of Jurkat T-cell membranes confirms that the localisation rate remains stable over 50 minute timeframes. **b)** WGA-HMSiR imaging of Jurkat T cell membranes similarly shows stable localisation over >1 hour timescales. **c)** anti-hCD45Fab imaging of a membrane protein is stable over 6 minutes.

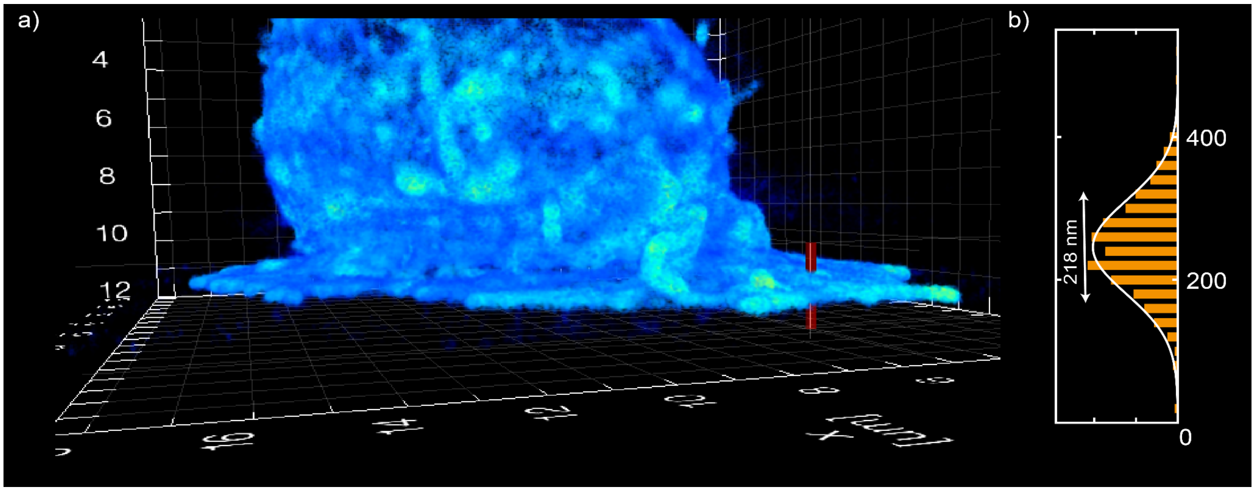


**Supplementary Figure 5: Thickness of the adhered Jurkat T cell membrane measured with resPAINT. a)** A line profile was applied to the skirt of the whole-cell image in Fig. 2g. **b)** A histogram of z position, shows the skirt FWHM thickness to be 218 nm.

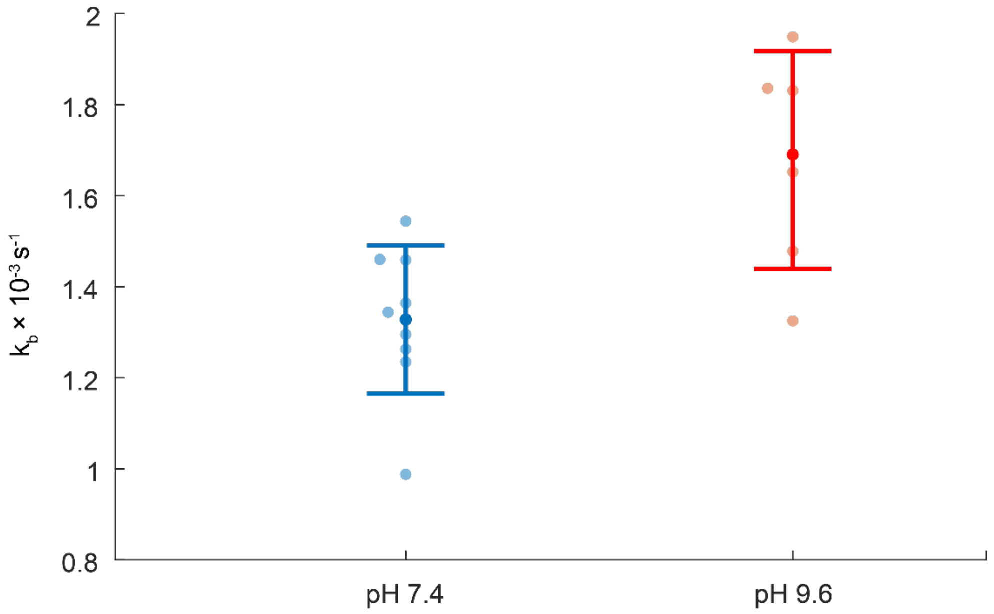


**Supplementary Figure 6: Off rates for anti-hCD45Fab-HMSiR on fixed cells.** a) The off rate, $k_b$, of the Fab fragment used in resPAINT of the membrane protein CD45 is measured on fixed cells by fitting the decay of fluorescence signal in a field of view to an exponential function at pH 7.4 and pH 9.6. The mean $k_b$ determined to be $1.33\times10^{-3}$ $s^{-1}$ at pH 7.4 and $1.68\times10^{-3}$ $s^{-1}$ at pH 9.6

a)

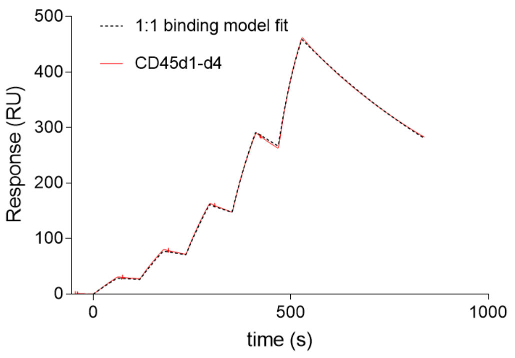


b)

| Maximum concentration CD45d1-d4 | $k_a$ ($M^{-1}$ $s^{-1}$) | $k_b$ ($s^{-1}$) | $K_D$ (nM) |
|---|---|---|---|
| 250 nM | 4.61e+4 | 1.59e-3 | 34.5 |
| 1000 nM | 5.48e+4 | 1.89e-3 | 34.6 |

**Supplementary Figure 7: Single cycle kinetic analysis of Gap8.3 binding to CD45d1-d4, using SPR at 20°C.** a) representative plot of Gap8.3 binding two-fold serially diluted CD45d1-d4 with a maximum concentration of 250 nM (red) and 1:1 binding model fit (dotted black). b) Summary table of kinetic constants derived from 1:1 binding model, where the $k_b$ was determined as $1.59\times10^{-3}$ $s^{-1}$.

**Supplementary Movie 1: PAINT *vs.* resPAINT**

We PAINT the apical surface of Jurkat T cell membranes using WGA, a lectin which binds to N-glycosyl moieties. The abundance of these targets renders WGA a popular membrane stain. For similar levels of background, resPAINT, with WGA-PAJF$_{549}$ (100 nM, photoactivation power density: 0.6 Wcm$^{-2}$), shows significantly increased localisation rate *vs.* conventional PAINT using WGA-AF$_{555}$ (100 pM). The exposure time is 30 ms.

**Supplementary Movie 2: resPAINT optimisation**

The effect of concentration and photoactivation power density is explored to determine optimal resPAINT conditions. The localisation rate increases with concentration and photoactivation power density, however so does the background levels. In the optimal condition (100 nM and 0.6 W cm$^{-2}$ laser power - highlighted at 14 s, background < 200 photons pixel$^{-1}$) the localisation rate averages ~1.5 loc frame$^{-1}$ at the apical cell surface. The exposure time is 30 ms.

**Supplementary Movie 3: Whole-cell resPAINT imaging**

Video representation of the whole Jurkat T cell membrane image from Fig 2c, highlighting the topographical features. Localisations are coloured by density as in Figure 2c. Grid spacing = 1 μm.

**Supplementary Movie 4: Cell-surface interaction**

Video representation of the whole Jurkat T cell membrane image from Fig 2g, highlighting the topographical features of the cell that are perturbed by electrostatic interactions with a PLL-coated glass surface. Localisations are coloured by depth as in Figure 2g. Grid spacing = 1 μm.

**Supplementary Movie 5: resPAINT with HMSiR**

Comparison of WGA-SiR, WGA-HMSiR at pH 7.4 and WGA-HMSiR at pH 9.6. Compared to SiR, HMSiR at pH 7.4 shows moderate localisation rate improvement. At the optimized pH of 9.6, the localisation rate is increased further, affording a 50-fold improvement compared to SiR. The exposure times is 20 ms.

**Supplementary Movie 6: Long-term resPAINT**

At an optimised probe concentration (0.1 nM), WGA-HMSiR imaging achieves a favorable localisation rate for DHPSF imaging (~1.77 loc. frame$^{-1}$). Three different timeframes are shown at the beginning (0 s), middle (2000 s) and end (4000 s) of a resPAINT experiment, demonstrating how the rate remains stable for at least 4000 s. The exposure time is 20 ms.

**Supplementary Movie 7: Optimal HMSiR pH**

Comparison of WGA-HMSiR at pH 9.6 with WGA-HMSiR at pH 11.5. At the same concentration of probe, WGA-HMSiR at pH 11.5 shows fewer and dimmer puncta. Under the same laser power and exposure time (20 ms), the pH-dominated duty cycle is faster than the exposure time, thus reducing the amount of collected photons.

**Supplementary Movie 8: Tetrapod PSF resPAINT**

WGA-SiR compared to WGA-HMSiR in similar background conditions using the tetrapod PSF. WGA-SiR shows few localisations for the same level of background. Meanwhile, due to the large footprint of the PSF, there is significant overlap between localisations. The exposure time was 30 ms.

**Supplementary Movie 9: Light field resPAINT**

As for Supplementary Video 8, but with SMLFM. Similarly, due to the large DOF, there is significant overlap between localisations. Single emitters are split into nine perspective views, where a minimum of three views are required to fit a localization. The exposure time was 20 ms.

**Supplementary Movie 10: Optimised Tetrapod PSF resPAINT**

An example of WGA-HMSiR imaging of a Jurkat T-cell membrane with the tetrapod PSF. The PSFs do not significantly overlap at this concentration (1 nM), and the background is minimal, despite the 10 µm DOF. The exposure time was 20 ms.

**Supplementary Movie 11: Optimised light field resPAINT**

An example of WGA-HMSiR imaging of a Jurkat T-cell membrane with the SMLFM. The PSFs do not significantly overlap at this concentration (5 nM), and the background is minimal, despite the 10 µm DOF. The exposure time was 20 ms.

**Supplementary Movie 12: resPAINT with a Fab**

αCD45 Gap8.3 Fab-SiR imaging is compared to αCD45 Gap8.3 Fab-HMSiR imaging. With HMSiR, the localisation rate is feasible and allows imaging of the membrane protein CD45. A negative control experiment using murine T cells shows a significantly lower rate, corresponding to a chance coincidence of ~9 %. The exposure time was 20 ms.

**Supplementary Movie 13: resPAINT simulation**

Simulation of 2D PAINT experiment where WGA is binding to the cell membrane. For PAINT, 1 nM WGA is simulated for HILO and LLS excitation, while for resPAINT, 100 nM photoactivatable WGA (0.001 $s^{-1}$ activation rate) is simulated. resPAINT greatly increases the localisation rate, while maintaining low background fluorescence.

**Supplementary Movie 14: Fab resPAINT simulation**

Simulation of 2D PAINT experiment where a Fab is binding to the cell membrane. For PAINT, 10 nM WGA is simulated for HILO and LLS excitation, while for resPAINT, 1000 nM photoactivatable WGA (0.001 $s^{-1}$ activation rate) is simulated. resPAINT greatly increases the localisation rate, while maintaining low background fluorescence.
